# Towards reliable named entity recognition in the biomedical domain

**DOI:** 10.1101/526244

**Authors:** John M. Giorgi, Gary D. Bader

## Abstract

**Motivation:** Automatic biomedical named entity recognition (BioNER) is a key task in biomedical information extraction (IE). For some time, state-of-the-art BioNER has been dominated by machine learning methods, particularly conditional random fields (CRFs), with a recent focus on deep learning. However, recent work has suggested that the high performance of CRFs for BioNER may not generalize to corpora other than the one it was trained on. In our analysis, we find that a popular deep learning-based approach to BioNER, known as bidirectional long short-term memory network-conditional random field (BiLSTM-CRF), is correspondingly poor at generalizing – often dramatically overfitting the corpus it was trained on. To address this, we evaluate three modifications of BiLSTM-CRF for BioNER to alleviate overfitting and improve generalization: improved regularization via variational dropout, transfer learning, and multi-task learning.

**Results:** We measure the effect that each strategy has when training/testing on the same corpus (“in-corpus” performance) and when training on one corpus and evaluating on another (“out-of-corpus” performance), our measure of the model’s ability to generalize. We found that variational dropout improves out-of-corpus performance by an average of 4.62%, transfer learning by 6.48% and multi-task learning by 8.42%. The maximal increase we identified combines multi-task learning and variational dropout, which boosts out-of-corpus performance by 10.75%. Furthermore, we make available a new open-source tool, called Saber, that implements our best BioNER models.

**Availability:** Source code for our biomedical IE tool is available at https://github.com/BaderLab/saber. Corpora and other resources used in this study are available at https://github.com/BaderLab/Towards-reliable-BioNER.

## 1 Introduction

PubMed contains over 30 million publications and is growing rapidly (https://www.nlm.nih.gov/bsd/index_stats_comp.html). Accurate, automated text mining tools are needed to maximize discovery and unlock structured information from this massive volume of text (Cohen and Hunter, 2008; Rzhetsky *et al.*, 2009).

A fundamental task in biomedical text mining, and text mining in general, is named entity recognition (NER). Biomedical named entity recognition (BioNER) is the task of identifying biomedical named entities — such as genes and gene products, diseases, cell types, chemicals, and species — in raw text. Biomedical named entities have several characteristics that make their recognition in text particularly challenging (Zhou *et al.*, 2004), including the use of descriptive entity names (e.g. “normal thymic epithelial cells”) leading to ambiguous term boundaries, and several spelling forms for the same entity (e.g. “N-acetylcysteine”, “N-acetyl-cysteine”, and “NAcetylCysteine”). Finding solutions for reliable BioNER has been (Kim *et al.*, 2004), and continues to be (Delėger *et al.*, 2016), a focus of various shared tasks organized within the biomedical natural language processing (BioNLP) community.

Recently, state-of-the-art methods have employed a domain-independent approach to this problem, based on deep learning and statistical word embeddings, called bidirectional long short-term memory network-conditional random field [BiLSTM-CRF (Lample *et al.*, 2016; Ma and Hovy, 2016)]. This approach has achieved state-of-the-art results for the task of BioNER (Habibi *et al.*, 2017). It is entity-agnostic, requiring only pre-trained word embeddings and labelled training data. Even more recently, improvements to this method have been made using transfer learning (Giorgi and Bader, 2018) and multi-task learning (Wang *et al.*, 2018), two forms of inductive transfer.

While BiLSTM-CRF paired with pre-trained word embeddings as inputs is a powerful approach which has found success in many sequence labelling tasks (Huang *et al.*, 2015), it contains a large number of trainable parameters which could lead to overfitting on the training data. For our purposes, we say that the model (i.e. BiLSTM-CRF) has overfit a dataset (i.e. a corpus of text annotated for biomedical entities) when it learns parameters that perform well on the training corpus but which do not generalize to other corpora annotated for the same biomedical entity. Overfitting on biomedical corpora appears to be a problem even for less-parameterized machine learning methods. For example, Gimli, an open-source tool for BioNER based on a CRF classifier, achieved a *F*_1_-score of 87.17% when trained and tested on the GENETAG corpus (Campos *et al.*, 2013a), but only a 45-55% *F*_1_-score when trained on the GENETAG (Tanabe *et al.*, 2005) corpus and tested on the CRAFT corpus (Bada *et al.*, 2012) for genes and proteins (Campos *et al.*, 2013b).

Galea *et al.* (2018) explore the issue of overfitting further, by demonstrating that performance of a CRF model for BioNER trained on individual corpora decreases substantially for recognition of the same biomedical entity in independent corpora. They found that overfitting may be partly caused by bias in popular BioNER evaluation corpora. In a simple orthographic feature analysis (i.e., what does a word look like?), they find that features which significantly predict biomedical named entities in one corpus (e.g., number of digits, number of capital letters, length of the text span) do not necessarily predict for the same entity in a different corpus. A powerful model such as BiLSTM-CRF is likely to overfit in this case, by learning representations that encode biases in individual corpora but which do not generalize well across corpora and to biomedical text at large. Indeed, in our analysis, we find that BiLSTM-CRF generalizes poorly to corpora other than those it was trained on, even when a relaxed scoring criterion is used. This problem is exacerbated by the fact that corpora in the biomedical domain tend to be small, and that many solutions for BioNER are trained and tested on the same corpora, e.g. the GENETAG corpus (Tanabe *et al.*, 2005) for gene and gene products, of which a modified version was used in the BioCreative II gene mention task (Smith *et al.*, 2008) and is sometimes referred to as the BC2GM corpus.

If BiLSTM-CRF models are to be truly useful in the large-scale annotation of widely diverse articles, such as the articles found in databases like PubMed, this overfitting problem will need to be addressed. To address this challenge, we sample the best ideas from recent work on BiLSTM-CRF models for BioNER (Crichton *et al.*, 2017; Habibi *et al.*, 2017; Giorgi and Bader, 2018; Wang *et al.*, 2018) and sequence labeling with BiLSTM-CRFs in general (Reimers and Gurevych, 2017) to propose several strategies to reduce overfitting and improve generalization, namely: additional regularization via variational dropout, transfer learning, and multi-task learning. We assessed the performance of the model on the same corpus it was trained on, which we call “in-corpus” performance, and when trained on one corpus and tested on another corpus annotated for the same entity class, which we call “out-of-corpus” performance. We used the latter as a measure of the model’s ability to generalize. We found that variational dropout improved out-of-corpus performance for 23 of 24 train/test corpus pairs, by an average of 4.62%, transfer learning 22 out of 24 train/test corpus pairs by an average of 6.48% and multi-task learning 24 out of 24 train/partner/test corpus pairs by an average of 8.42%. All of these strategies achieved an improvement in out-of-corpus performance without degrading the average in-corpus performance. We also found that certain combinations of these strategies lead to additive boosts in performance. The best combination was multi-task learning and variational dropout, which together boost out-of-corpus performance by 10.75%. Finally, we make available to the community a user-friendly, open-source tool for BioNLP (“Saber”) which incorporates these best practices: https://github.com/BaderLab/saber.

## 2 Materials and methods

The following sections present a technical explanation of the basic neural network architectures used in this study. We first briefly describe LSTM, a specific kind of recurrent neural network (RNN), before introducing the architecture of the BiLSTM-CRF model for sequence labelling tasks (Lample *et al.*, 2016) used in this study and in prior work (Habibi *et al.*, 2017; Giorgi and Bader, 2018; Wang *et al.*, 2018). We then describe several proposed modifications to this model’s architecture and training strategies aimed at improving generalization. Finally, we describe the corpora used for evaluation, training details, and evaluation metrics.

### 2.1 Bidirectional Long Short-Term Memory-Conditional Random Field (BiLSTM-CRF)

LSTMs, a type of RNN architecture, are a popular choice for sequence labelling tasks due to their ability to use previous information in a sequence for processing of current input. An LSTM achieves this behaviour through the use of a memory cell, which serves as a summary of the preceding elements of an input sequence, and is able to model dependencies between sequence elements even if they are far apart (Hochreiter and Schmidhuber, 1997). The input to an LSTM unit is a sequence of vectors of length *T*, 
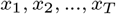
for which it produces an output sequence of vectors of equal length, 
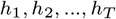
by applying a non-linear transformation learned during the training phase. Each *h*_*t*_ is called the activation of the LSTM at token *t*, where a token is an instance of a sequence of characters in a document that are grouped together as a useful semantic unit for processing. The formula to compute one activation of an LSTM unit in the LSTM-CRF model is provided below (Lample *et al.*, 2016): 
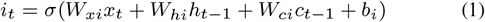

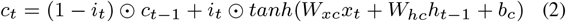

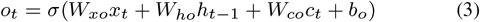

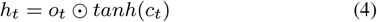
where all *W* s and *b*s are trainable parameters, *σ*(·) and *tanh*(·) denote the element-wise sigmoid and hyperbolic tangent activation functions, and ⊙ is the element-wise product.

Intuitively, an LSTM unit is able to “remember” information about previous items in a sequence (such as words in a sentence) by modifying the cell state (*c*_*t*_). It regulates the flow of information in the cell state through a series of gates (called the “update” and “forget” gates), which themselves are simple neural network layers who’s behaviour is learned during the training process. Additionally, a final “output” gate decides what information from the cell state to use when processing the next item in the sequence.

The above-described LSTM-layer processes input in one direction, and thus can only encode dependencies on elements that came earlier in the sequence. Processing information *after* the current element is also useful. Thus, another LSTM-layer which processes input in the reverse direction is commonly used. The resulting network is called a bidirectional LSTM [BiLSTM (Graves and Schmidhuber, 2005)]. The representation of a word using this model is obtained by concatenating its left and right context representations, 

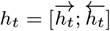
which effectively encodes a word in its context.

A simple sequence labelling approach would be to use the *h*_*t*_’s as features to make independent tagging decisions for each output *y*_*t*_. For example, one simple strategy would be to select the label *y*_*i*_ that has the highest probability in *h*_*i*_, i.e., 
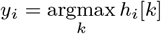
where *k* is the number of distinct tags. However, this greedy approach fails to consider the dependencies between subsequent labels. Therefore, instead of modelling tagging decisions independently, BiLSTM-CRF models them jointly using a conditional random field [CRF (Lafferty *et al.*, 2001)].

Each input sequence token is mapped to a pre-trained word embedding, which is concatenated with a learned character representation of that token (see Section 2.2). This character-enhanced word embedding is processed by a BiLSTM network, before being fed to a dense layer which maps the outputs of this BiLSTM network to a sequence of vectors containing the probability of each label for each corresponding token. Finally, a CRF is used to output the most likely sequence of predicted labels based on these probabilities.

Figure 1 illustrates the basic BiLSTM-CRF architecture. With the exception of the pre-trained word embeddings, all layers of the network are learned jointly. A detailed description of the architecture is explained in Lample *et al.* (2016). A similar architecture, which uses a convolutional neural network (CNN) in place of an LSTM to learn character-level representations, is presented in Ma and Hovy (2016).

**Fig. 1.**
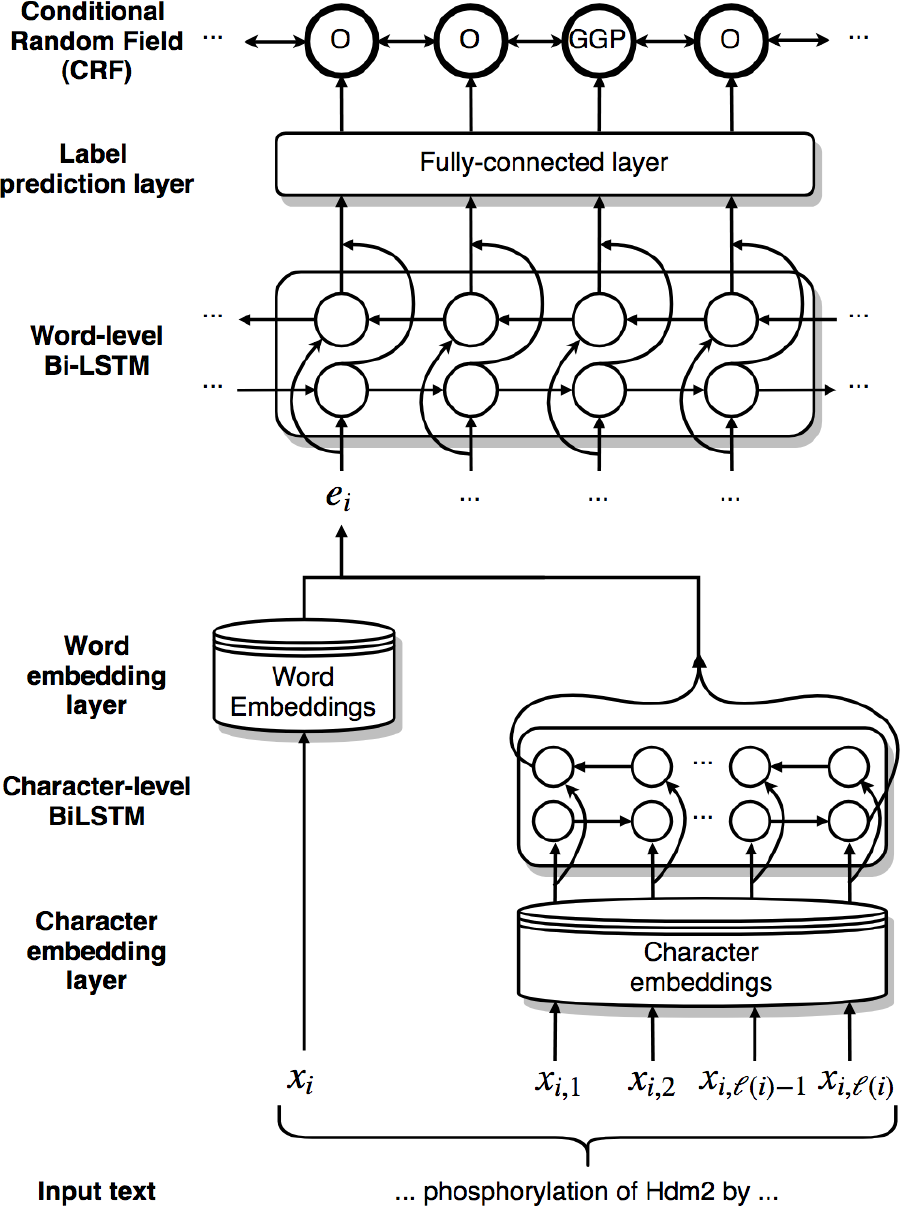
Architecture of the bidirectional long short-term memory network-conditional random field (BiLSTM-CRF) model for named entity recognition (NER). Here, *x*_*i*_ is the *i*^*th*^ token in the input sequence, *x*_*ij*_ is the *j*^*th*^ character of the *i*^*th*^ token, *ℓ*(*i*) is the number of characters in the *i*^*th*^ token and *e*_*i*_ is the character-enhanced token embedding of the *i*^*th*^ token. Output labels “O” represent tokens “outside” of an entity, and label “GGP” stands for “gene or gene product”. Figure from Giorgi and Bader (2018).

### 2.2 Word embeddings

The inputs to the BiLSTM-CRF model for sequence labelling are the individual words (or tokens) of a given sentence. A standard approach is to utilize statistical word embedding techniques which capture functional (i.e., semantic and syntactic) similarity of words based on their surrounding words. Such word embeddings are learned on large unlabelled datasets, typically containing billions of words (Collobert *et al.*, 2011; Mikolov *et al.*, 2013; Pennington *et al.*, 2014). The learned continuous vector representations, or word embeddings, encode many linguistic patterns. In the canonical example, the resulting vector for vec(“king”) *−* vec(“man”) + vec(“woman”) is closest to the vector associated with “queen”, i.e., vec(“queen”). These token-based word embeddings effectively capture distributional similarities of words (where does the word tend to occur in a corpus?). To capture orthographic similarities (what does the word look like?) and better handle out-of-vocabulary tokens and misspellings, character-based word representation models (Ling *et al.*, 2015) have been developed. By using each individual character of a token to generate the token vector representation, character-based word embeddings encode sub-token patterns such as morphemes (e.g. suffixes and prefixes), morphological inflections (e.g. number and tense) and other information not contained in the token-based word embeddings. The BiLSTM-CRF architecture used in this study combines character-based word representations with token-based word representations to produce character-enhanced word representations, allowing the model to learn both distributional and orthographic features of words.

In this study, we used the *Wiki-PubMed-PMC* model, trained on a combination of PubMed abstracts (nearly 23 million abstracts) and PubMedCentral (PMC) articles (nearly 700,000 full-text articles) plus approximately four million English Wikipedia articles, therefore mixing domain-specific texts with domain-independent ones. The model was created by Pyysalo *et al.* (2013) using Google’s word2vec tool (Mikolov *et al.*, 2013). We chose this model because previous work has shown it to work well for the task of BioNER (Habibi *et al.*, 2017; Giorgi and Bader, 2018; Wang *et al.*, 2018).

### 2.3 Variational Dropout

Given the over-parameterization of neural networks, generalization performance crucially relies on the ability to regularize a model sufficiently. Dropout is a technique to prevent neural networks from overfitting by reducing the amount of co-adaptation between units in a neural network (Srivastava *et al.*, 2014). The basic idea is to randomly discard or “drop” units from the network during training. Previous applications of BiLSTM-CRF models for BioNER (Habibi *et al.*, 2017; Giorgi and Bader, 2018) have relied on a single dropout layer applied to the character-enhanced word-embeddings, the final inputs to the word-level BiLSTM layers. This strategy was proposed by Lample *et al.* (2016) as a means of encouraging the model to make use of both character- and word-derived information in the embedding. However, no regularization technique was applied to the recurrent layers of the model containing the majority of the model’s trainable parameters, which were, therefore, free to overfit on the training data. The seemingly simple solution to this problem is to apply dropout to the recurrent layers of the model. Unfortunately, regularizing recurrent connections with naive applications of dropout is ineffective, as it disrupts the RNNs ability to retain long-term dependencies (Pachitariu and Sahani, 2013; Bayer *et al.*, 2013; Zaremba *et al.*, 2014).

Recently, Gal and Ghahramani (2016) have proposed a solution to this problem, which they call variational dropout, where the same units are dropped across multiple time steps (in our case, tokens of an input sentence) as opposed to randomly dropping units at each time step (Figure 2). It was demonstrated that applying variational dropout to the input, recurrent, and output connections of an RNN significantly outperforms no dropout and naive applications of dropout. We hypothesize that regularizing the recurrent layers of the BiLSTM-CRF model via variational dropout will greatly reduce the degree to which the model overfits the training corpus, especially in the case where the training corpus is small. To test this, we compare the performance of two BiLSTM-CRF models for the task of BioNER: one with the dropout strategy proposed by Lample *et al.* (2016) and used in previous work (Habibi *et al.*, 2017; Giorgi and Bader, 2018), and a second model in which variational dropout is additionally applied to the input, recurrent, and output connections of all recurrent layers.

**Fig. 2.**
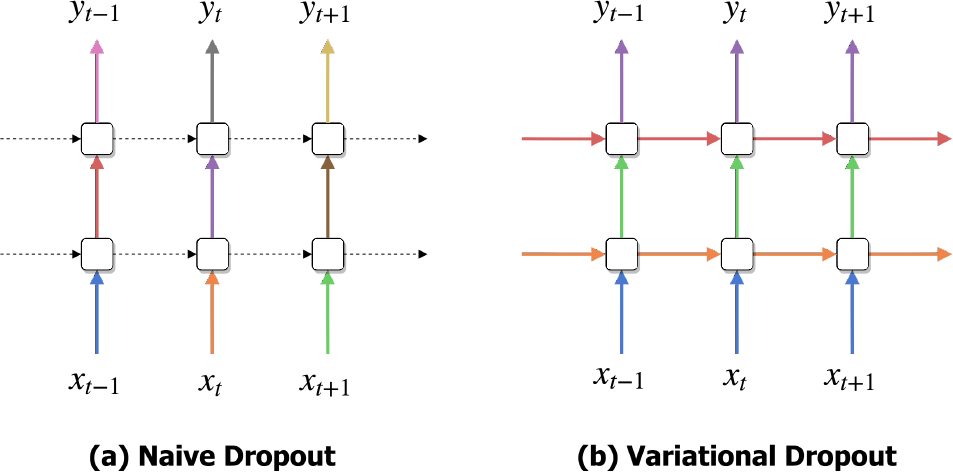
Naive dropout in recurrent neural networks (RNNs) compared to variational dropout, as proposed by Gal and Ghahramani (2016). Each square represents the unit of an RNN, horizontal arrows the recurrent connections, and vertical arrows the input and output connections. Coloured arrows represent connections to which dropout has been applied, with individual colours corresponding to different dropout masks. Dashed lines represent standard connections with no dropout. In naive dropout (a), a different dropout mask is randomly sampled for each time step and, typically, no dropout is applied to the recurrent layers of the neural network. In variational dropout (b), the same dropout mask is used across all time steps for input, recurrent and output connections. Figure adapted from Gal and Ghahramani (2016).

### 2.4 Transfer Learning

Transfer learning is a machine learning research problem which aims to perform a task on a “target” dataset using knowledge learned from a “source” dataset (Pan and Yang, 2010; Li, 2012; Weiss *et al.*, 2016). A formal definition of transfer learning involves defining a domain 𝒟 and a task 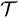. A domain consists of a feature space *χ* and a marginal probability distribution *P* (*X*) over the feature space, where *X* = *x*_1_, …, ∈ χ is the set of all input features. Situations where the domain differs between tasks are typically referred to as domain adaptation (Bridle and Cox, 1991; Ben-David *et al.*, 2010). For a given domain, 𝒟 = {*χ*, *P* (*X*)}, a task 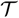 consists of a label space 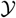, and a conditional probability distribution *P* (*Y*|*X*), where 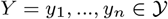 is the set of all targets. The goal of transfer learning is to learn the target conditional probability distribution *P* (*Y*_*t*_|*X*_*t*_) for the target task 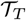 with the information gained from the source task 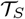, where 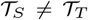. In our setting, the feature space *χ* contains the character-enhanced word embeddings, the final representation of words fed to the LSTM layers, and the label space 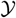 for a task 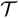 contains the set of biomedical entity classes (e.g. gene/protein) annotated in the training dataset for that task 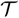.

For neural networks, transfer learning is typically implemented by using some or all of the learned parameters of a neural network pre-trained on a source dataset to initialize training for a second neural network to be trained on a target dataset. Ideally, transfer learning improves generalization of the model, reduces training times on the target dataset, and reduces the amount of labelled data needed to obtain high performance. Recently, we demonstrated that a simple transfer learning strategy for BioNER (Giorgi and Bader, 2018) reduces the amount of labelled data needed to achieve high performance from a BiLSTM-CRF model. However, our previous work did not assess the impact of using transfer learning to improve generalizability.

In this study, we explore the effect of transfer learning on the generalizability of a BiLSTM-CRF model for BioNER. We apply transfer learning in a near identical way to Giorgi and Bader (2018), by first training on a large, silver-standard corpus (SSC), i.e., the source task 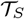, and then using the learned parameters to initialize training on a smaller, gold-standard corpus (GSC), i.e., the target task 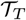. GSCs are manually annotated by biological experts. As such, they tend to be small, but highly reliable. In contrast, SSCs are annotated in an automatic or semi-automatic fashion (e.g., by deploying existing NER tools on a large body of unlabelled text) and therefore tend to be much larger than GSCs, but of much lower quality. Like Giorgi and Bader (2018), we use CALBC (Collaborative Annotation of a Large Biomedical Corpus) as our silver-standard corpus (Rebholz-Schuhmann *et al.*, 2010), specifically CALBC-SSC-III (Kafkas *et al.*, 2012). The motivation for using transfer learning here is that by exposing the network to a much larger number of examples, it may learn representations that apply more generally to the task of BioNER, instead of learning representations that encode biases in the individual training corpora.

### 2.5 Multi-task learning

Multi-task learning (Caruana, 1993) is a machine learning method in which multiple learning tasks are solved at the same time. In the classification context, multi-task learning is used to improve the performance of multiple classification tasks by learning them jointly. The idea is that by sharing representations between tasks, we can exploit commonalities, leading to improved learning efficiency, prediction accuracy, and generalizability for the task-specific models, when compared to training the models separately (Thrun, 1996; Caruana, 1998; Baxter *et al.*, 2000). Multi-task learning has been used successfully for many applications of machine learning, including natural language processing (Collobert and Weston, 2008), speech recognition (Deng *et al.*, 2013), computer vision (Girshick, 2015) and drug discovery (Ramsundar *et al.*, 2015).

Recent work has explored multi-task learning for the task of BioNER. Crichton *et al.* (2017) demonstrated that a neural network multi-task model outperforms a comparable single-task model, on average, for the task of BioNER. Similarly, Wang *et al.* (2018) also found that multi-task learning outperformed single-task learning for BioNER with a BiLSTM-CRF. However, neither study explored the effect of multi-task learning on the model’s ability to generalize. Given that, similar to transfer learning, multi-task learning exposes the model to a much larger number of training examples, we hypothesize that multi-task learning will improve generalization. Additionally, because multi-task learning typically requires the optimization of multiple objective or loss functions, we hypothesize that it may serve as a form of regularization for the BiLSTM-CRF model and thus reduce overfitting, especially on small training corpora. To test this, we compare the performance of a single-task and multi-task BiLSTM-CRF model for the task of BioNER.

**Table 1.** The gold-standard corpora (GSCs) used in this work.

| Entity type | Corpus | No. annotations |
| --- | --- | --- |
| Chemicals | BC4CHEMD (Krallinger *et al.*, 2015) | 84,310 |
|  | BC5CDR (Li *et al.*, 2016) | 15,935 |
|  | CRAFT (Bada *et al.*, 2012) | 6053 |
| Diseases | BC5CDR | 12,852 |
|  | NCBI-Disease (Doğan *et al.*, 2014) | 6881 |
|  | Variome (Verspoor *et al.*, 2013) | 5859 |
| Species | CRAFT | 6868 |
|  | Linnaeus (Gerner *et al.*, 2010) | 4263 |
|  | S800 (Pafilis *et al.*, 2013) | 3708 |
| Genes/proteins | BC2GM (Smith *et al.*, 2008) | 24,583 |
|  | CRAFT | 16,114 |
|  | JNLPBA (Kim *et al.*, 2004) | 35,336 |

For our baseline (BL) single-task model, we use the same architecture introduced by Lample *et al.* (2016) and described here in Section 2.1. Specifically, we use a BiLSTM-CRF to jointly model the word and character sequences in a given input sentence (Figure 1). Our multi-task model (MTM) builds off this BiLSTM-CRF architecture. The MTM is a global model comprised of distinct, task-specific input and output layers, while the hidden layers (and their parameters) are shared across all tasks (Figure 3). While it is possible to construct different configurations of the MTM (i.e. by sharing or not sharing particular layers between tasks) we follow Wang *et al.* (2018) who found it best to share all hidden layers of the BiLSTM-CRF model for BioNER. In our setting, each dataset the model is trained on represents a single task, and we therefore use the terms *task* and *dataset* interchangeably. A multi-task model (MTM), therefore, consists of *m* different models, each trained on a separate dataset, *D*_*m*_, while sharing some of the model parameters across datasets. During training, the model optimizes the log-probability of the correct tag sequence for each dataset. In practice, the model is trained in an end-to-end fashion.

**Fig. 3.**
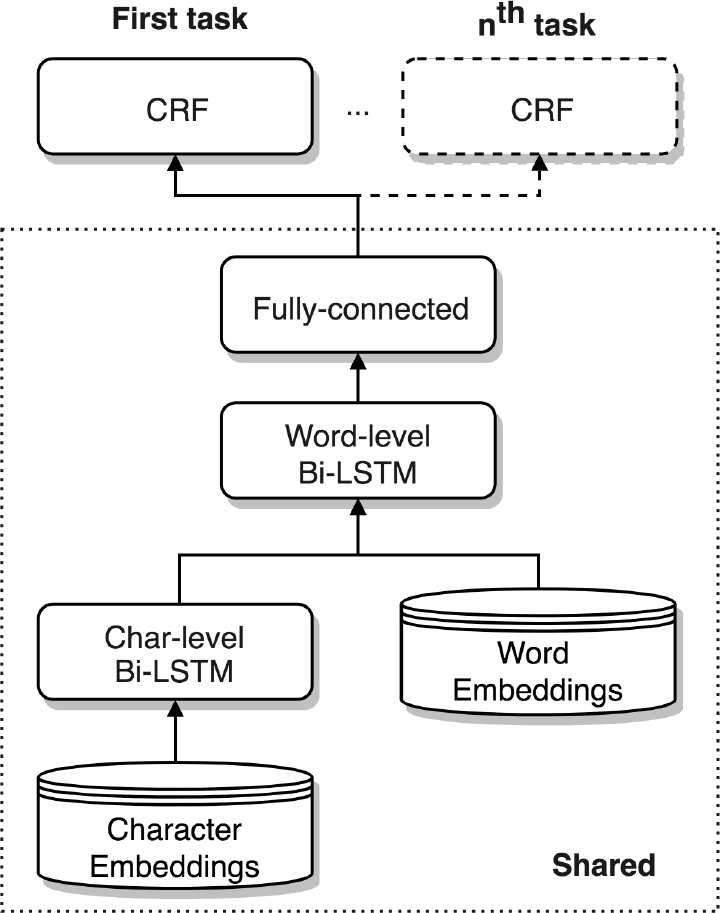
Architecture of the bidirectional long short-term memory network-conditional random field (BiLSTM-CRF) multi-task model (MTM). For each task, there is a distinct input layer (not shown), which connects to the shared hidden layers. At the output, there are distinct conditional random field (CRF) layers corresponding to each of the input layers or tasks. The model is capable of supporting an arbitrary number of input datasets. In our setting, each dataset the model is trained on represents a single task, and we therefore use the terms ‘task’ and ‘dataset’ interchangeably

### 2.6 Datasets

We used a total of 12 biomedical GSCs for our experiments, each containing hand-labelled annotations for one or more of the major biomedical entity classes: chemicals, diseases, species, and genes/proteins. In our transfer learning experiments, we use a random 100,000 abstracts from the silver-standard CALBC-III-Small corpus (Kafkas *et al.*, 2012) as our source dataset. Containing over 2 million annotations, this corpus subset is many times larger than any of the GSCs.

One complication in the transfer learning experiments is that some of the abstracts in the SSC are also present in the GSC, thus we removed such cases using a blacklist of duplicated PubMed IDs. We also compiled a blacklist of single-token entities that appear in the SSC, and are present in at least one of the GSCs but never annotated (see Supplementary Material). These are mostly higher level entity categories, like “Genes”, “elderly”, “infertile” and “epitope”, and arguably do not constitute biomedical named entities. We found this de-noising step to be important for good performance during transfer learning (Giorgi and Bader, 2018).

Word labels for each dataset are encoded using the BIO tag scheme. In this scheme, a word describing a protein or gene entity, for example, is tagged with “B-PRGE” if it is at the beginning of the entity and “I-PRGE” if it is in the middle of the entity. All other words that do not describe any specific entities are tagged as “O”.

Nearly all of the GSCs were originally collected by Crichton *et al.* (2017), and are publicly available at https://github.com/cambridgeltl/MTL-Bioinformatics-2016. We evaluated our models on especially popular corpora, in order to compare our results to other studies which evaluate deep learning methods for BioNER (Crichton *et al.*, 2017; Habibi *et al.*, 2017; Giorgi and Bader, 2018; Wang *et al.*, 2018). Table 1 lists each corpora along with its size in number of annotations.

### 2.7 Hyperparameters and model training

The selection of good hyperparameters for a given neural network is a time-consuming, but necessary task (Snoek *et al.*, 2012). Optimal hyperparameters can often make the difference between mediocre and state-of-the-art performance; commonly tuned hyperparameters include the learning rate, batch size, and dropout rate. In spite of this, previous applications of BiLSTM-CRF to BioNER have been presented without a strong validation for why certain hyperparameter values were chosen (Habibi *et al.*, 2017; Giorgi and Bader, 2018; Wang *et al.*, 2018), which is likely due to the computational cost of determining optimal hyperparameters via methods such as grid search, randomized search or more advanced optimization techniques such as Tree-structured Parzen Estimator [TPE (Bergstra *et al.*, 2011)]. Recently, Reimers and Gurevych (2017) presented a set of empirically validated, optimal hyperparameters for deep BiLSTM networks for sequence labelling tasks, which we follow in this work (Table 2). Some hyperparameters, such as the dropout rate, were modified across experiments and are presented in their respective Results sections.

**Table 2.** Hyperparameter values and model details of the bidirectional long short-term memory network-conditional random field (BiLSTM-CRF) model used in this study.

| Hyperparameter | Value | Comment |
| --- | --- | --- |
| Word embeddings | Wiki-PubMed-PMC | Wiki-PubMed-PMC model (Pyysalo *et al.*, 2013), trained on a combination of PubMed abstracts, PubMedCentral (PMC) articles and approximately four million English Wikipedia articles using Google’s word2vec (Mikolov *et al.*, 2013). |
| Character representation | LSTM | Character representations in the BiLSTM-CRF model can be learned either with convolutional neural networks (CNNs) (Ma and Hovy, 2016) or long short-term memory (LSTM) networks (Lample *et al.*, 2016). |
| Optimizer | Nadam | Parameters follow those provided in Dozat (2016). |
| Gradient normalization | $\Gamma = 1$ | A common strategy to deal with the exploding gradient problem (Pascanu *et al.*, 2013). Rescales the gradient whenever the norm goes over some threshold $\Gamma$. |
| Tagging scheme | BIO | In the BIO tagging scheme, each token of the training corpus is tagged as occurring at the beginning of an entity mention (B), inside an entity mention (I) or outside an entity mention (O). |
| Dropout rate | 0.3 | Fraction of units to drop in a given layer. Dropout was applied to the character-enhanced word embeddings. |
| No. LSTM layers | 2 | The number of word-level LSTM layers. |
| No. Recurrent units | 100 | The number of units in each LSTM layer within the BiLSTM-CRF model. Because all LSTM layers are bi-directional in this model, the total number of units per LSTM layer is 200. |
| Mini-batch size | 32 | When using a batch size $\neq 1$, input sequences must be padded. We truncated or right-padded every input sequence in the training corpus to match a length of 100 tokens. Similarly, every character sequence was truncated or right-padded to match a length of 25 characters. |

In all experiments, the BiLSTM-CRF model was trained for 50 epochs, using the back-propagation algorithm to update the parameters on every training step. In each training session, the character embedding vectors are initialized randomly and learned jointly with the other parameters of the model. The word embedding vectors are retrieved directly from a pre-trained word embedding lookup table and the classical Viterbi algorithm is used to infer the final labels for the CRF model. The model was implemented using Keras (https://keras.io/).

In the transfer learning setting, a BiLSTM-CRF model was first trained on the CALBC-SSC-III corpus for a single epoch. The learned weights were then used to initialize training on one of the 12 GSCs introduced in Section 2.6. The weights of the final layer of the model, the CRF, are randomly initialized and the state of the optimizer is reset when a model is transferred.

When training the MTM, each dataset is used to update the parameters of the model, in random order. For example, if the multi-task model is trained on two datasets, we first randomly select one of these datasets and use it to update all shared parameters, and parameters specific to that dataset (via forward and backward propagation), followed by using the remaining dataset to update all shared parameters, and parameters specific to that dataset. We consider this process a single epoch. The model thus behaves like separate neural networks, each training on its own dataset and performing prediction with its own output layer.

### 2.8 Evaluation metrics

For each experiment, we measure the performance of the model in terms of how it performs on the same corpus it was trained on, which we refer to as “in-corpus” performance, as well as how it performs on a corpus other than the one it was trained on, or “out-of-corpus” performance – sometimes called *translational* or *cross-corpus* performance. We use out-of-corpus performance to measure the model’s generalizability. When evaluating in-corpus performance, we used five-fold cross-validation, and report the average performance from across the five folds. When evaluating out-of-corpus performance, we used an entire corpus for training – except for 10% of examples which are used as a validation set – and a second, entire corpus for testing. Each out-of-corpus experiment was run three times and average results are reported.

We compared all methods in terms of *F*_1_-score on the test sets. *F*_1_-score is computed as the harmonic mean of precision (P) and recall (R), where precision is the percentage of predicted labels that are gold labels (i.e., labels that appear in the training corpus), and recall is the percentage of gold labels that are correctly predicted: 
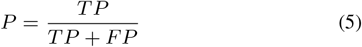

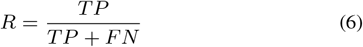

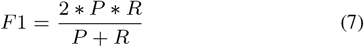
where TP is the number of true positives predicted by the model, FP the number of false positives and FN the number of false negatives. In the case of in-corpus performance, a predicted label was considered correct if and only if it exactly matched a gold label. Because annotation standards differ across corpora, we used a relaxed matching criterion when evaluating out-of-corpus performance, namely, right-boundary matching, which was found to be a suitable alternative to exact boundary matching in an evaluation of multiple BioNER matching strategies by Tsai *et al.* (2006). In this scheme, a predicted label is considered correct as long as the right boundary of the text span matched the gold label (e.g., if the gold label is “familial gastric cancer” and the predicted label “gastric cancer”, the prediction is scored as a true positive).

## 3 Results

### 3.1 Establishing a baseline

To establish a strong baseline for each corpus used in this study, we performed five-fold cross-validation using the hyperparameters presented in Section 2.7. We use the dropout strategy proposed by Lample *et al.* (2016) and employed by Habibi *et al.* (2017) and Giorgi and Bader (2018), which applies a single dropout layer to the final inputs to the word-level BiLSTM layer, the character-enhanced word embeddings, with a dropout rate of 0.3. The hope is that by randomly dropping dimensions from the character-enhanced word embeddings, the model will learn to make use of both character-derived and word-derived information in the embedding. In Table 3, we present our baseline *F*_1_-scores along with the best reported *F*_1_-scores from Habibi *et al.* (2017), Wang *et al.* (2018), and Giorgi and Bader (2018), all of whom use nearly identical BiLSTM-CRF models and the same word embeddings as used in this study, and Crichton *et al.* (2017), who used a CNN-based model for BioNER.

**Table 3.**
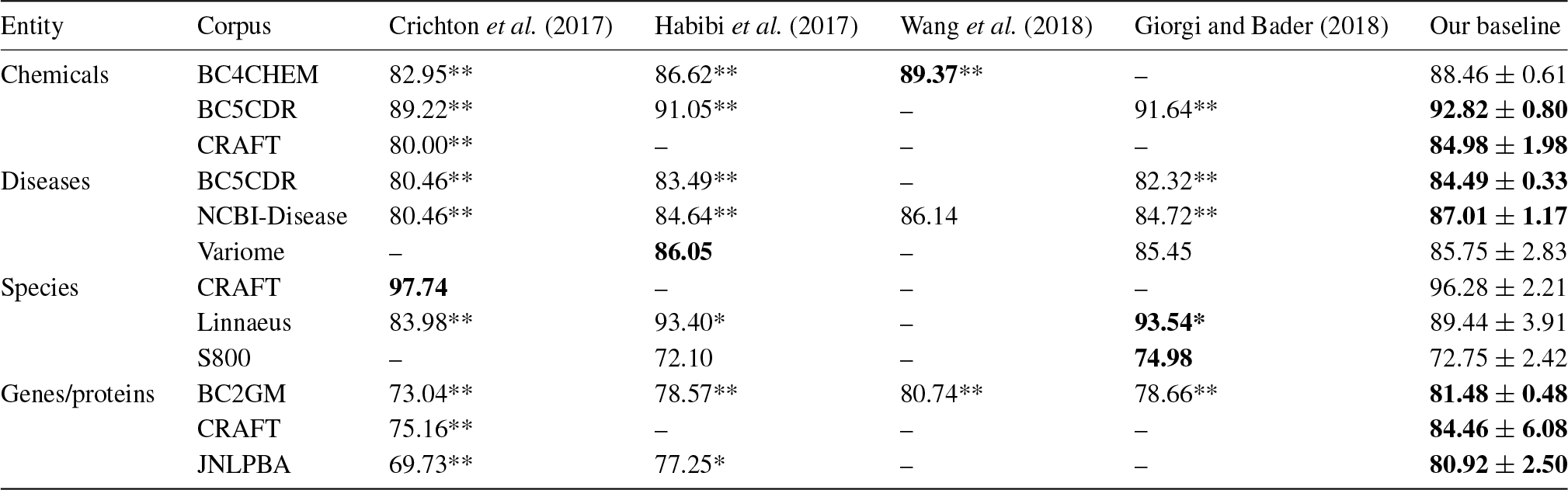
Previously published state-of-the-art *F*_1_-scores for biomedical named entity recognition (BioNER) for all corpora used in this study. All scores are compared to the scores obtained by our baseline model via five-fold cross-validation, using an exact matching criteria. Statistical significance is measured through a two-tailed t-test. Bold: best scores, *: significantly different than our baseline (*p* ≤ 0.05), **: significantly different than our baseline (*p* ≤ 0.01).

Our baseline significantly outperforms previous results obtained with BiLSTM-CRF models for seven out of the twelve corpora evaluated and was comparable to the best method in the remaining five cases. Since our architecture is nearly-identical and paired with the same word embeddings, this is likely due to our choice of optimal hyperparameters as presented by Reimers and Gurevych (2017). For the remainder of the study, we compare all performance scores to our baseline.

### 3.2 Can BiLSTM-CRF generalize for BioNER?

To illustrate the poor generalizability of a BiLSTM-CRF model for BioNER, we trained the model on various corpora and evaluated its performance on independent corpora annotated for the same entity type. Even when corpora are annotated for the same entity type, they are unlikely to capture the same underlying distribution because they were both in-corpus and out-of-corpus performance. In Table 5, we compare the in-corpus performance of the baseline (BL) BiLSTM-CRF model employing a simple dropout strategy – proposed by Lample *et al.* (2016) and used in previous applications of BiLSTM-CRF to BioNER (Habibi *et al.* (2017); Giorgi and Bader (2018)) – compared to a model in which variational dropout (VD) has been additionally applied to the recurrent layers. We use dropout ratios of 0.3, 0.3 and 0.1 for the input, output and recurrent connections, respectively. In Table 6, we measure the effect that made by different groups with no explicit consensus about what should be annotated. To control for the expected drop in performance due to differing annotation guidelines, we used a relaxed matching criterion in our evaluation (described in Section 2.8). We found that, even with a relaxed matching criterion, performance of the model as measured by *F*_1_-score falls by an average of 31.16% when the model is evaluated on a corpus other than the one it was trained on (Table 4). These results demonstrate the dramatically poor generalizability of BiLSTM-CRF for BioNER, even though the model obtains state-of-the-art results when trained and tested on the same corpus (Table 3). Out-of-corpus performance was worst for chemicals, falling by an average of 33.45%, followed by genes/proteins (29.27%), then by species (28.47%), and disease (26.09%). This ranking largely appeals to intuition. Gene and protein nomenclature, for example, is highly irregular. In contrast, the naming of species of living things follows a formal, standard naming system (i.e. binomial nomenclature, following the International Code of Zoological Nomenclature, or ICZN). In a single case — when the model was trained on BC4CHEMD and tested on BC5CDR — out-of-corpus performance is better than in-corpus performance. This suggests that the annotations in BC5CDR are well-predicted by a model trained on BC4CHEMD, and therefore the relaxed matching criterion used when evaluating out-of-corpus but not in-corpus performance offers an unfair advantage.

**Table 4.** In-corpus (IC) and out-of-corpus (OOC) performance, measured by *F*_1_-score, of the bidirectional long short-term memory-conditional random field model (BiLSTM-CRF). IC performance is derived from five-fold cross-validation, using an exact matching criteria. OOC performance is derived by training on one corpus (train) and testing on another annotated for the same entity type (test) using a relaxed, right-boundary matching criteria.

| Entity | Train | Test | IC | OOC | $\Delta F_1$ |
| --- | --- | --- | --- | --- | --- |
| Chemicals | BC4CHEMD | BC5CDR | 88.46 | 90.90 | -2.43 |
|  |  | CRAFT | — | 47.44 | 41.02 |
|  | BC5CDR | BC4CHEMD | 92.82 | 71.81 | 21.00 |
|  |  | CRAFT | — | 39.55 | 53.27 |
|  | CRAFT | BC4CHEMD | 84.98 | 40.50 | 44.47 |
|  |  | BC5CDR | — | 41.59 | 43.39 |
| Diseases | BC5CDR | NCBI-Dis. | 84.49 | 76.67 | 7.82 |
|  |  | Variome | — | 74.03 | 10.46 |
|  | NCBI-Dis. | BC5CDR | 87.01 | 69.62 | 17.38 |
|  |  | Variome | — | 74.98 | 12.03 |
|  | Variome | BC5CDR | 85.75 | 22.45 | 63.30 |
|  |  | NCBI-Dis. | — | 40.17 | 45.58 |
| Species | CRAFT | Linnaeus | 96.28 | 45.32 | 50.96 |
|  |  | S800 | — | 36.88 | 59.40 |
|  | Linnaeus | CRAFT | 89.44 | 82.49 | 6.95 |
|  |  | S800 | — | 62.90 | 26.54 |
|  | S800 | CRAFT | 72.75 | 57.09 | 15.65 |
|  |  | Linnaeus | — | 61.43 | 11.31 |
| Genes/proteins | BC2GM | CRAFT | 81.48 | 56.04 | 25.44 |
|  |  | JNLPBA | — | 69.77 | 11.72 |
|  | CRAFT | BC2GM | 84.46 | 44.11 | 40.35 |
|  |  | JNLPBA | — | 52.88 | 31.58 |
|  | JNLPBA | BC2GM | 80.92 | 51.03 | 29.88 |
|  |  | CRAFT | — | 44.29 | 36.62 |

**Table 5.** In-corpus (IC) performance, measured by *F*_1_-score, of the baseline (BL) bidirectional long short-term memory-conditional random field (BiLSTM-CRF) compared to a BiLSTM-CRF with variational dropout (VD). In the BL model, dropout is applied only to the character-enhanced word embeddings. In the VD model, dropout is additionally applied to the input, recurrent, and output connections of all LSTM layers. IC performance is derived from five-fold cross-validation, using an exact matching criteria. Statistical significance is measured through a two-tailed t-test. Bold: best scores, *σ*: standard deviation, *: significantly different than the BL (*p* ≤ 0.05), **: significantly different than the BL (*p* ≤ 0.01).

| Entity | Corpus | BL |  | VD |  |
| --- | --- | --- | --- | --- | --- |
| | | Avg. | $\sigma$ | Avg. | $\sigma$ |
| Chemicals | BC4CHEMD | 88.46 | 0.61 | **88.71** | 0.76 |
|  | BC5CDR | 92.82 | 0.80 | **93.08** | 0.82 |
|  | CRAFT | 84.98 | 1.98 | **85.22** | 1.37 |
| Disease | BC5CDR | 84.49 | 0.33 | **85.10** | 0.56 |
|  | NCBI-Dis. | 87.01 | 1.17 | **87.60** | 1.50 |
|  | Variome | **85.75** | 2.83 | 85.69 | 3.81 |
| Species | CRAFT | 96.28 | 2.21 | **96.38** | 2.26 |
|  | Linnaeus | 89.44 | 3.91 | **89.66** | 7.47 |
|  | S800 | 72.75 | 2.42 | **77.39** | 4.17 |
| Genes/proteins | BC2GM | 81.48 | 0.48 | **83.10\*\*** | 0.50 |
|  | CRAFT | 84.46 | 6.08 | **86.09** | 5.19 |
|  | JNLPBA | 80.92 | 2.50 | **81.95** | 2.62 |

**Table 6.**
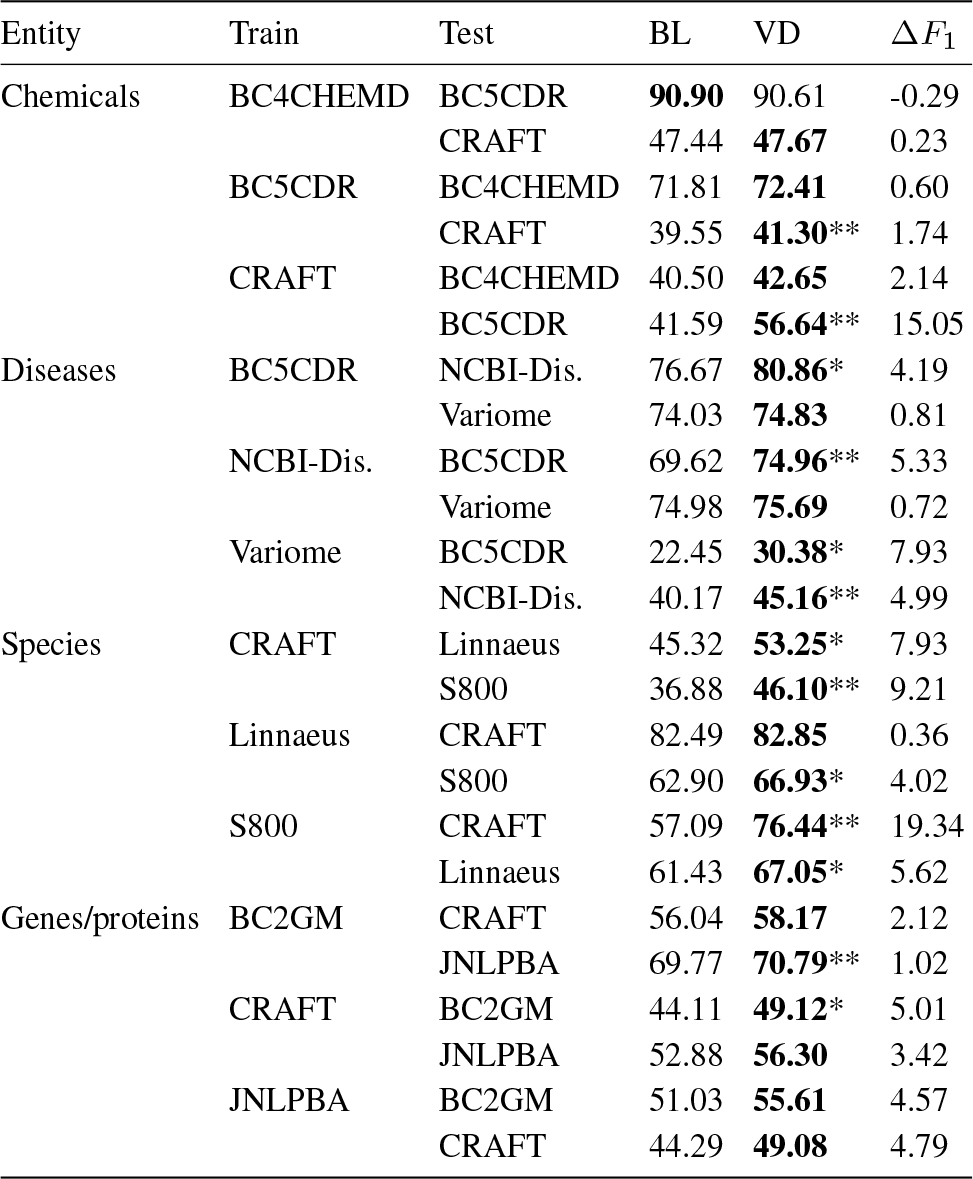
Out-of-corpus (OOC) performance, measured by *F*_1_-score, of the baseline (BL) bidirectional long short-term memory-conditional random field (BiLSTM-CRF) compared to a BiLSTM-CRF with variational dropout (VD). In the BL model, dropout is applied only to the character-enhanced word embeddings. In the VD model, dropout is additionally applied to the input, recurrent, and output connections of all LSTM layers. OOC performance is derived by training on one corpus (train) and testing on another annotated for the same entity type (test) using a relaxed, right-boundary matching criteria. Bold: best scores, *: significantly different than the BL (*p* ≤ 0.05), **: significantly different than the BL (*p* ≤ 0.01).

### 3.3 Improved regularization

In this experiment, we explore the effect that additional regularization of the BiLSTM-CRF model via variational dropout (see Section 2.3) has on variational dropout has on the generalizability of the model, by comparing out-of-corpus performance to the baseline.

In general, variational dropout has a small positive impact on in-corpus performance. For at least two corpora, S800 and BC2GM, variational dropout leads to a large improvement in performance, but only the latter case is statistically significant. For out-of-corpus performance, variational dropout improves performance for nearly every train/test corpus pair we evaluated, with an average improvement of 4.62%. In some cases, variational dropout leads to sizable improvements in out-of-corpus performance, such as when the model was trained on S800 and tested on CRAFT (19.34%), or trained on CRAFT and tested on BC5CDR (15.05%). In one case - when trained on BC4CHEMD and tested on BC5CDR - variational dropout reduced out-of-corpus performance, although the performance difference was minimal (less than 0.5%). This corroborates our hypothesis in Section 3.2 that BC5CDR is well-predicted by BC4CHEMD, and may explain why additional regularization failed to improve performance in this case.

In summary, variational dropout improves the out-of-corpus performance of the model, without degrading in-corpus performance. Regularization of the recurrent layers of a BiLSTM-CRF model via variational dropout is, therefore, a straightforward way to improve the generalizability of the model for BioNER.

### 3.4 Transfer learning

In this experiment, we quantify the effect that a transfer learning strategy for BiLSTM-CRF has on both in-corpus and out-of-corpus performance. In Table 7, we compare the in-corpus performance of the baseline BiLSTM-CRF model (BL) to that of the model trained with transfer learning (TL) and in Table 8 we similarly compare out-of-corpus performance of the two models. In the transfer learning setting, the model was pre-trained on the CALBC-SSC-III, which is annotated for chemicals, diseases, species and genes/proteins, before being trained on one of the 12 GSCs, each annotated for a single entity class. The state of the optimizer is reset during this transfer, but model weights for all layers beside the final CRF layer are retained (See Sections 2.4 and 2.7).

**Table 7.** In-corpus (IC) performance, measured by *F*_1_-score, of the baseline (BL) bidirectional long short-term memory-conditional random field (BiLSTM-CRF) compared to a BiLSTM-CRF trained with transfer learning (TL). The TL model was pre-trained on the CALBC-Small-III corpus. IC performance is derived from five-fold cross-validation, using an exact matching criteria. Statistical significance is measured through a two-tailed t-test. Bold: best scores, *σ*: standard deviation, *: significantly different than the BL (*p* ≤ 0.05), **: significantly different than the BL (*p* ≤ 0.01).

| Entity | Corpus | BL |  | TL |  |
| --- | --- | --- | --- | --- | --- |
| | | Avg. | $\sigma$ | Avg. | $\sigma$ |
| Chemicals | BC4CHEMD | 88.46 | 0.61 | **88.98** | 0.63 |
|  | BC5CDR | **92.82** | 0.80 | 92.20 | 0.86 |
|  | CRAFT | 84.98 | 1.98 | **85.80** | 1.74 |
| Disease | BC5CDR | **84.49** | 0.33 | 84.41 | 0.24 |
|  | NCBI-Dis. | 87.01 | 1.17 | **87.66** | 0.86 |
|  | Variome | 85.75 | 2.83 | **86.69** | 3.03 |
| Species | CRAFT | 96.28 | 2.21 | **96.55** | 1.72 |
|  | Linnaeus | 89.44 | 3.91 | **90.72** | 4.90 |
|  | S800 | 72.75 | 2.42 | **74.93** | 3.27 |
| Genes/proteins | BC2GM | **81.48** | 0.48 | 80.65* | 0.57 |
|  | CRAFT | 84.46 | 6.08 | **85.50** | 4.59 |
|  | JNLPBA | 80.92 | 2.50 | **81.56** | 2.73 |

**Table 8.**
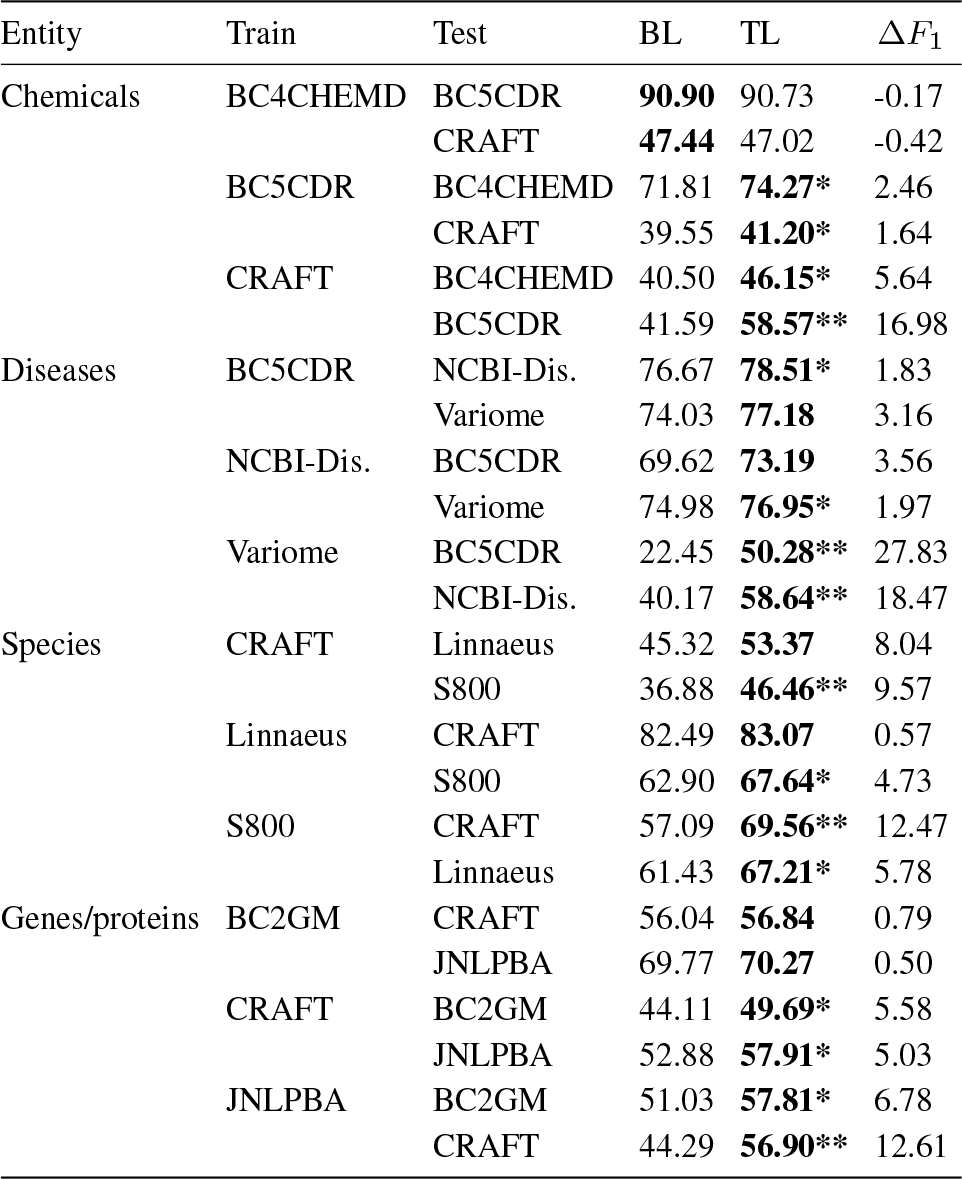
Out-of-corpus (OOC) performance, measured by *F*_1_-score, of the baseline (BL) bidirectional long short-term memory-conditional random field (BiLSTM-CRF) compared to a BiLSTM-CRF trained with transfer learning (TL). The TL model was pre-trained on the CALBC-Small-III corpus. OOC performance is derived by training on one corpus (train) and testing on another annotated for the same entity type (test) using a relaxed, right-boundary matching criteria. Statistical significance is measured through a two-tailed t-test. Bold: best scores, *: significantly different than the BL (*p* ≤ 0.05), **: significantly different than the BL (*p* ≤ 0.01).

In general, transfer learning had a small positive effect on in-corpus performance, boosting the average *F*_1_-score by approximately 1%. In contrast, transfer learning had a large positive effect on out-of-corpus performance, improving performance for nearly every train/test pair we evaluated for an average improvement of 6.48%. In a handful of cases, such as when the model was trained on the CRAFT corpus and tested on the BC5CDR corpus, performance improved by over 10%. In a single case (i.e. when the model was trained on Variome and tested on BC5CDR) transfer learning doubled out-of-corpus performance over the baseline.

We conclude that the use of transfer learning improves the generalizability of BiLSTM-CRF models for BioNER, in some cases dramatically, while preserving in-corpus performance.

### 3.5 Multi-task learning

To assess the effect that a multi-task learning strategy for BiLSTM-CRF has on both in-corpus and out-of-corpus performance, we evaluate a model trained on all corpus pairs within an entity class. For each model, there is a set of “train”, “partner” and “test” corpora. We define “train” to be the corpus a model was trained on, “partner” to be the second corpus included in the training session under the MTM, and “test” to be the corpus the model was evaluated on. In Table 9, we compare the in-corpus performance of the single-task, baseline BiLSTM-CRF model, to that of the multi-task model (MTM). In Table 10 we similarly compare out-of-corpus performance of the BL and MTM. In our experiments, multi-task learning is implemented nearly identically to Wang *et al.* (2018): by training a BiLSTM-CRF model - which shares the parameters of all hidden layers - on more than one dataset simultaneously (See Sections 2.5 and 2.7).

**Table 9.**
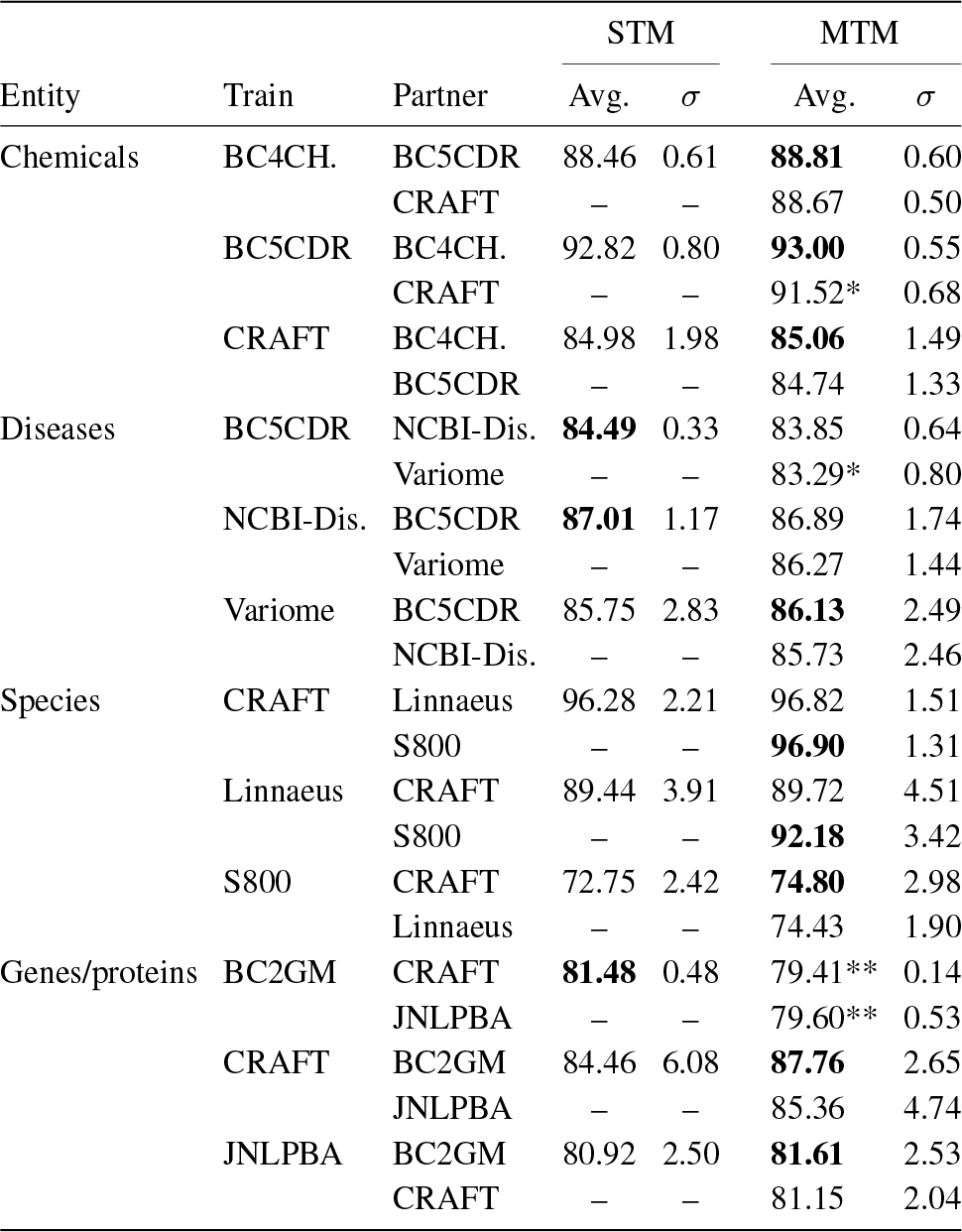
In-corpus (IC) performance, measured by *F*_1_-score, of the baseline (BL) bidirectional long short-term memory-conditional random field (BiLSTM-CRF) compared to the multi-task model (MTM). The multi-task model (MTM) is trained on pairs of corpora (train, partner), where each corpus is used during training to update the parameters of all hidden layers. IC performance is derived from five-fold cross-validation, using an exact matching criteria. Statistical significance is measured through a two-tailed t-test. Bold: best scores, *σ*: standard deviation, *: significantly different than the BL (*p* ≤ 0.05), **: significantly different than the BL (*p* ≤ 0.01).

**Table 10.**
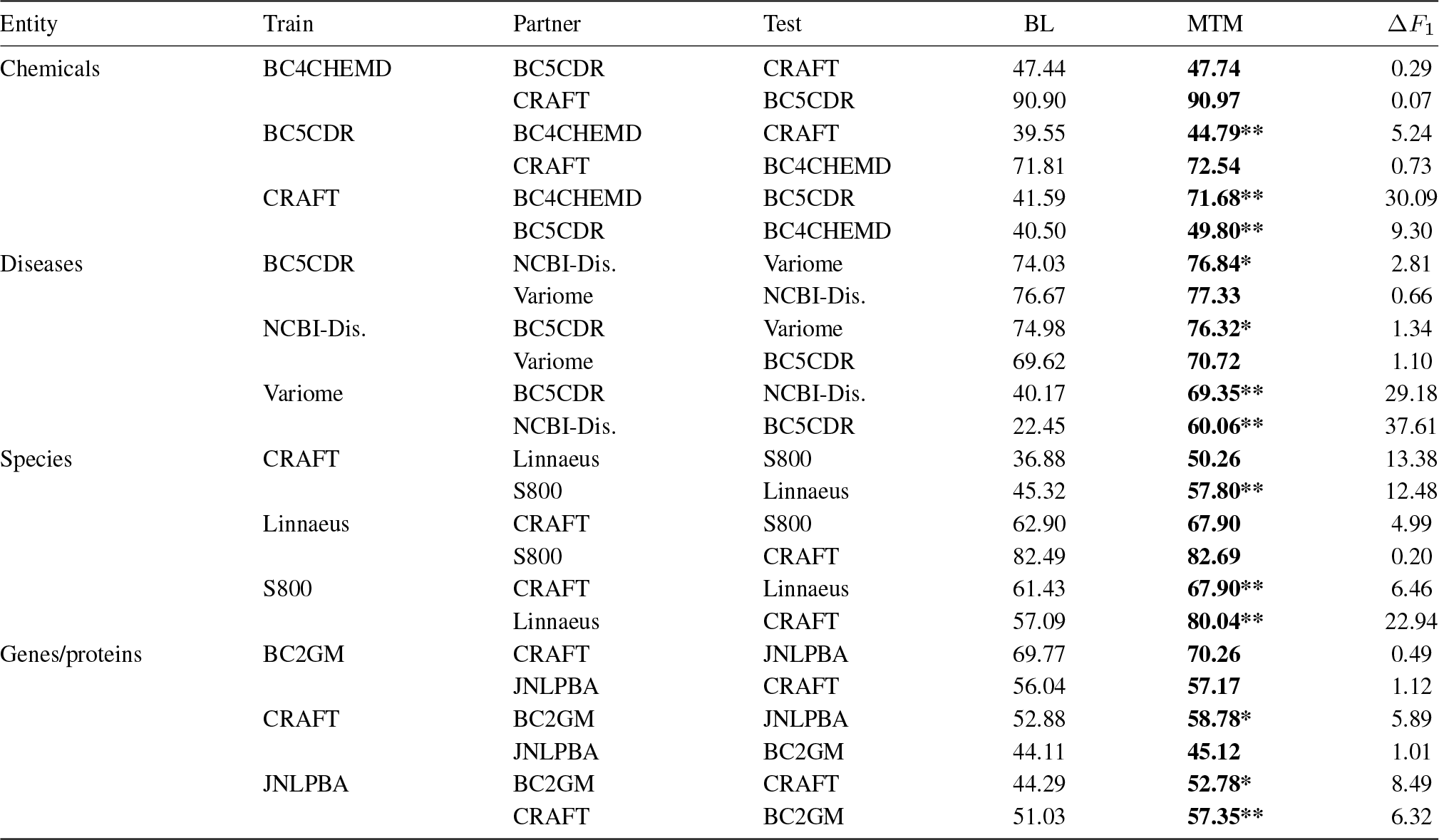
Out-of-corpus (OOC) performance, measured by *F*_1_-score, of the baseline (BL) bidirectional long short-term memory-conditional random field (BiLSTM-CRF) compared to the multi-task model (MTM). The multi-task model (MTM) is trained on pairs of corpora (train, partner), where each corpus is used during training to update the parameters of all hidden layers. OOC performance is derived by training on a pair of corpora (train, test) and testing on another corpus annotated for the same entity type (test) using a relaxed, right-boundary matching criteria. Bold: best scores, *: significantly different than the BL (*p* ≤ 0.05), **: significantly different than the BL (*p* ≤ 0.01).

Multi-task learning, as applied here, appears to have little impact on in-corpus performance. In a few cases, such as when the model was trained on BC2GM alongside CRAFT or JNLPBA, the MTM significantly unperformed the baseline. Despite this, average performance of the BL and the MTM were nearly identical, at 85.74% and 85.99% respectively. Multi-task learning improved out-of-corpus performance for every train/partner/test corpus pair we evaluated, with an average improvement of 8.42% (Table 10). In some cases, this improvement was substantial, such as when the model was trained on the Variome and NCBI-Disease corpora and tested on the BC5CDR corpus (37.61%). However, we do observe significant variability overall in the degree of improvement, suggesting that multi-task learning is sensitive to the choice of train/partner pairs.

In summary, a multi-task learning strategy of simultaneously training on multiple datasets appears to be an effective way to significantly boost out-of-corpus performance of BiLSTM-CRF models for BioNER. Our results do suggest, however, that both in-corpus and out-of-corpus performance of the multi-task model are sensitive to the choice of partner corpus.

### 3.6 Combining the proposed modifications

We next evaluate if combinations of the proposed modifications improve BiLSTM-CRFs model performance above individual modifications (Figure 4). In general, all combinations of the proposed modifications improve average out-of-corpus performance without degrading in-corpus performance. However, not all combinations are additive. For example, multi-task learning improves out-of-corpus performance by 8.42%, transfer learning by 6.48% but together only by 5.61%. The biggest boost to out-of-corpus performance is achieved by the MTM paired with additional regularization of the recurrent layers via variational dropout, improving average performance by 10.75%. Therefore, we recommend this combination of strategies to produce a model with the highest expected out-of-corpus performance.

**Fig. 4.**
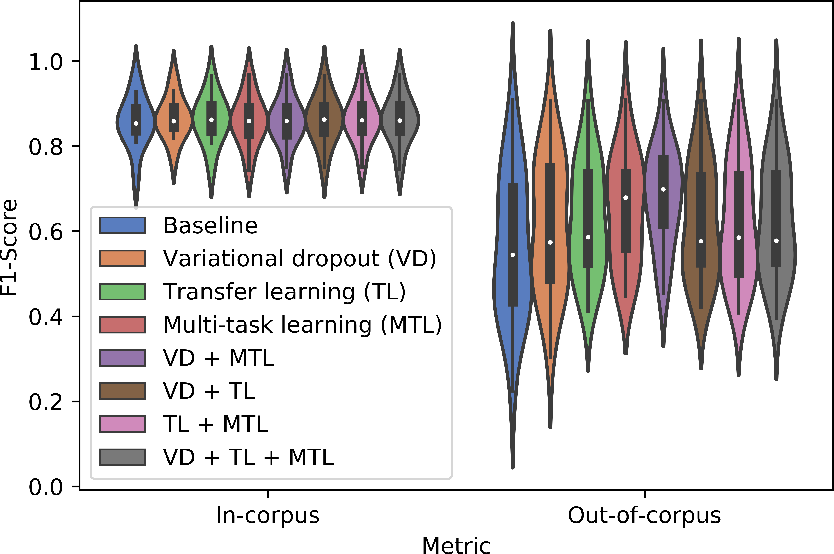
Violin plot of the average in-corpus (IC) and out-of-corpus (OOC) performance, measured by *F*_1_-score, of the bidirectional long short-term memory-conditional random field (BiLSTM-CRF) model. IC performance is derived from five-fold cross-validation, using an exact matching criteria. OOC performance is derived by training on one corpus (train) and testing on another corpus annotated for the same entity type (test) using a relaxed, right-boundary matching criterion. The average performance of a model employing one of each of the proposed modifications: variational dropout (VD), transfer learning (TL) and multi-task learning (MTL) independently as well as models which employ all combinations of these methods is shown.

## 4 Discussion

In this study, we demonstrated that BiLSTM-CRF, a popular deep learning-based approach to BioNER, does a poor job generalizing, often dramatically over-fitting the corpus it was trained on. We quantified the degree of over-fitting by first establishing a state-of-the-art baseline model, and then evaluated its performance when trained and tested on the same corpus (in-corpus performance) and when trained on one corpus and tested on another annotated for the same entity type (out-of-corpus performance). Using out-of-corpus performance to measure generalization, we found that even with a relaxed matching criterion, out-of-corpus performance of the model falls by an average of 31.16%. We then evaluated three modifications - variational dropout, transfer learning, and multi-task learning and demonstrated that these modifications improve generalizability and reduce over-fitting. On average, variational dropout improves out-of-corpus performance by 4.62%, transfer learning by 6.48% and multi-task learning by 8.42%. Importantly, each modification improves average out-of-corpus performance without degrading average in-corpus performance. Finally, we found that some combinations of these modifications lead to additive improvements in performance, most notably multi-task learning paired with variational dropout, which improved average out-of-corpus performance by 10.75%.

The modifications we propose in this work are often quite easy to implement in practice. For example, using a framework like Keras (https://keras.io/), implementing variational dropout is a matter of providing dropout rates as arguments to an LSTM layer (https://keras.io/layers/recurrent/#lstm). Sharing the parameters of hidden layers across models (i.e., the multi-task learning setting) is as easy as defining these layers globally and re-using them when defining a new model. We provide our model to the community as an easy-to-use tool for BioNLP under a permissive MIT license (https://github.com/BaderLab/saber).

We note three limitations of our work. Firstly, certain combinations of these modifications are not additive. For example, multi-task learning improves out-of-corpus performance by 8.42%, transfer learning by 6.48% but together only by 5.61%. Based on our experiments, we recommend using multi-task learning paired with variational dropout, which boosted out-of-corpus performance by 10.75%. Secondly, the most promising modification to BiLSTM-CRF in our experiments, multi-task learning, appears to be sensitive to the choice of train/partner corpus pairs. A user should therefore evaluate multiple train/partner sets to determine the most beneficial set of training corpora for their use case. Thirdly, while transfer learning is not part of our recommended combination, we previously found it to significantly improve performance on smaller datasets (less than 6000 labels) (Giorgi and Bader, 2018), which we did not evaluate here. Presumably, it would still be useful and should be considered in these cases.

### 4.1 Related Work

In our multi-task experiments, we restricted ourselves to training only on two corpora, which were annotated for the same entity type. These restrictions were made only to keep the number and the training time of the multi-task experiments within a reasonable range and not because of inherent limitations in the model’s architecture. Previous work has suggested that multi-task learning with deep neural networks for the task of BioNER may increase performance even when trained on corpora that do not annotate the same entity class. Crichton *et al.* (2017) report, for example, that the best partner for Linnaeus (Species) out of 15 datasets was NCBI-Disease (Disease), not another dataset which was annotated for Species. Additionally, previous work (Wang *et al.*, 2018) has suggested that training a BiLSTM-CRF on many corpora (i.e., greater than two) in a multi-task setting leads to sizable improvements in BioNER. Thus, a further direction for our work could be to explore performance improvements as increasing numbers of corpora annotated for different entity types are used for training. We suspect that this could significantly boost out-of-corpus and generalization performance. These findings, combined with our work here, provide an interesting new direction for deep-learning based approaches to BioNER, one that moves away from training/testing on individual corpora and towards one in which a single model is trained on many corpora (and even additionally pre-trained on an extremely large SSC), to produce a robust and reliable tagger suitable for deployment on massive literature databases (such as PubMed).

Our strategy for transfer learning involves pre-training a model on a large SSC and transferring the learned weights to initialize training on a smaller, but typically much higher quality, GSC. As we were writing this paper, a novel transfer learning strategy for NLP demonstrated state-of-the-art performance on many benchmark datasets (Radford *et al.*, 2018; Howard and Ruder, 2018; Devlin *et al.*, 2018). This transfer learning strategy involves first training a language model on a massive corpus of unlabeled text. The task of a language model is to predict the next most probable word given a sequence of words. By learning this task, the model is required to capture both syntax and semantics, and is also required to encode something akin to common sense. This is followed by the addition of task-specific layers, which take the output of the language model as input and are trained on labelled data in order for the model to learn some specific classification task such as NER. This transfer learning strategy has already been applied to BioNER with some success (Sachan *et al.*, 2018). In the future, we plan to explore this transfer learning strategy for BioNER, and also for other tasks in the biomedical text-mining pipeline, such as relation and event extraction.

## 5 Conclusion

While biomedical named entity recognition (BioNER) has recently made substantial advances in performance with the application of deep learning, current applications suffer from poor generalizability. In this study, we first quantified the out-of-corpus performance of a bidirectional long short-term memory network-conditional random field (BiLSTM-CRF) model for BioNER by training on one corpus and testing on another annotated for the same entity type, our measure of the model’s ability to generalize. We found that even with a relaxed scoring criteria, performance dropped by an average of 31.16% when compared to a state-of-the-art baseline. We then explored three modifications: variational dropout, transfer learning, and multi-task learning, which reduce the model’s tendency to overfit. Furthermore, we show that combining multi-task learning and variational dropout under a single model boosts out-of-corpus performance by over 10%. We propose that our model will significantly outperform previous applications of BiLSTM-CRF models to BioNER when deployed for the large-scale annotation of widely diverse articles, such as the articles found in databases like PubMed. We make our model accessible as an easy-to-use BioNLP tool, Saber (https://github.com/BaderLab/saber).

## Acknowledgements

This research was enabled in part by support provided by Compute Ontario (https://computeontario.ca/) and Compute Canada (www.computecanada.ca). We also gratefully acknowledge the support of NVIDIA Corporation with the donation of the Titan Xp GPU used for this research‥

