## Additional File 1 for "Towards reliable named entity recognition in the biomedical domain"

This document contains supplementary information for the paper *Towards reliable named entity recognition in the biomedical domain.*

### 1 Entity blacklists for silver-standard corpora

In an effort to reduce noise in the SSCs, a selection of single-token entities present but not annotated in any of the GSCs were removed from the SSCs. For example, certain text spans such as 'genes', 'elderly', and 'infertile' are annotated in the SSCs but not annotated in any of the GSCs, and so were removed from the SSCs. Entity blacklists are available in **additional_file_1.zip**. Alternatively, they are available at https://github.com/BaderLab/Towards-reliable-BioNER.

### 2 PMID blacklists for silver-standard corpora

In the transfer learning experiments of our paper, we first train on a large, automatically annotated silver standard corpus (SSC) before training on manually annotated gold-standard corpora (GSCs). To ensure that we weren't training and then testing on the same documents (i.e., a document present in the training set of the SSC is not present in the testing set of any of the GSCs), we simply removed any document present in SSC that was present in any of the GSCs. PMID blacklists are available in **additional_file_2.zip**. Alternatively, they are available at https://github.com/BaderLab/Towards-reliable-BioNER.
